# Vegetative nuclear positioning is required for calcium and ROS signaling in Arabidopsis pollen tubes

**DOI:** 10.1101/2020.02.10.942722

**Authors:** Morgan Moser, Andrew Kirkpatrick, Norman Reid Groves, Iris Meier

## Abstract

Efficient transport and delivery of sperm cells (SCs) is vital for angiosperm plant fertility. In *Arabidopsis thaliana*, SCs are transported through the growing pollen tube by a connection with the vegetative nucleus (VN). During pollen tube growth, the VN leads the way and maintains a fixed distance from the pollen tube tip, while the SCs lag behind the VN. Upon reception at the ovule, the pollen tube bursts and the SCs are released for fertilization. In pollen tubes of Arabidopsis mutants *wit12* and *wifi*, deficient in the outer nuclear membrane component of a plant LINC complex, the SCs precede the VN and the VN falls behind. Subsequently, pollen tubes frequently fail to burst upon reception. In this study, we sought to determine if the pollen tube reception defect observed in *wit12* and *wifi* is due to decreased sensitivity to reactive oxygen species (ROS). Here we show that *wit12* and *wifi* are hyposensitive to exogenous H_2_O_2_, and that this hyposensitivity is correlated with decreased proximity of the VN to the pollen tube tip. Additionally, we report the first instance of nuclear Ca^2+^ spikes in growing pollen tubes, which are disrupted in the *wit12* mutant. In the *wit12* mutant, nuclear Ca^2+^ spikes are reduced in response to exogenous ROS, but these spikes are not correlated with pollen tube burst. This study finds that VN proximity to the pollen tube tip is required for both response to exogenous ROS, as well as internal nuclear Ca^2+^ fluctuations.

**Summary:** Mutants deficient in outer nuclear membrane proteins display defects in reactive oxygen species-induced pollen tube burst and nuclear Ca^2+^ signatures that correlate with the position of the vegetative nucleus.

## Introduction

Angiosperm fertilization requires delivery of the non-motile sperm cells (SCs) to the ovules by pollen tubes. This process is guided by a variety of interactions between the pollen tube and the surrounding sporophytic and finally gametophytic female tissue. While significant progress has been made towards understanding the underlying male-female crosstalk and the signaling components involved, the final step, pollen tube reception and termination, still is among the least understood (Dresselhaus and Franklin-Tong, 2013; Higashiyama and Yang, 2017; Johnson et al., 2019).

Pollen tube reception at the ovule involves intricate interactions between the pollen tube and the synergid cells, leading to pollen tube growth arrest and burst, and the release of the SCs (Kessler and Grossniklaus, 2011; Dresselhaus and Franklin-Tong, 2013). Several signaling components of this step on the female side have been identified (For review see (Johnson et al., 2019)). Loss of function mutants of the synergid cell surface receptor kinase FERONIA lead to failed WT pollen tube burst upon arriving at a *fer* mutant ovule, and subsequent pollen tube overgrowth (Escobar-Restrepo et al., 2007). FERONIA is required for an increase in reactive oxygen species (ROS) production by the synergid cells upon pollen tube arrival and a synergid calcium (Ca^2+^) fluctuation signal that precedes a corresponding Ca^2+^ spike in the pollen tube tip prior to burst (Iwano et al., 2012; Ngo et al., 2014). *In vitro*-applied ROS leads to pollen tube growth arrest and rupture in a Ca^2+^-dependent manner, suggesting pollen tube-synergid crosstalk involves a synergid-triggered Ca^2+^-ROS signal that leads to pollen tube termination and sperm cell release (Duan et al., 2014).

How signals from the synergids are perceived by the pollen tube is less well understood. Two FERONIA-related proteins, ANXUR1 and ANXUR2, are specifically expressed in pollen, and the double mutant shows premature growth arrest and burst, but whether they are involved in signal reception by the pollen tube is not known (Boisson-Dernier et al., 2009). A second pair of pollen tube receptors (BUPS1 and BUPS2) has a similar function, protecting the pollen tube from premature burst. BUPS1 and BUPS2 bind both pollen-expressed and female-expressed Rapid Alkalinization Factor (RALF) peptide ligands, and interaction with female-expressed RALF34 releases the protection from burst by BUPS1 and BUPS2 (Ge et al., 2017). Thus, an interplay between ovule-derived and pollen-derived peptide ligands might enable the pollen tube to respond to the synergid environment created by FERONIA and related pathways (Ge et al., 2017)

During pollen germination and subsequent pollen tube growth, the two SCs are physically connected with the pollen tube nucleus (“vegetative nucleus”, VN), and collectively named the male germ unit (MGU) (Borg et al., 2009). As the pollen tube grows, the MGU maintains a fixed distance from the advancing pollen tube tip, with the VN leading the SCs (Li et al., 2018). The fact that the VN and SCs migrate as a unit has been proposed to be important for efficient movement of the SCs to the ovule (Russell and Cass, 1981; Dumas et al., 1985; McCue et al., 2011). A mutant that transports only the VN is phenotypically near-normal, including pollen tube rupture after entering the ovule (Zhang et al., 2017), suggesting that reception and termination signaling can proceed in the absence of the SCs.

In contrast, mutants that transport the SCs, but lead to a partial loss of the VN at the pollen tube tip have defects in pollen tube termination (Zhou and Meier, 2014). The mutations causing this effect are in genes coding for the plant outer nuclear membrane (ONM) proteins WIP (WIP1, WIP2, and WIP3) and WIT (WIT1 and WIT2), the components of a plant LINC complex (Zhou et al., 2012). LINC complexes are protein complexes spanning the inner nuclear membrane (INM) and ONM, thus connecting the nucleus to cytoplasmic components, such as the cytoskeleton (Starr and Fridolfsson, 2010). The core of LINC complexes is composed of the ONM Klarsicht/ANC-1/Syne homology (KASH) proteins, with interact with the INM Sad1 and UNC-84 (SUN) proteins in the nuclear envelope lumen. In animals, LINC complexes are often involved in nuclear movement and nuclear positioning (Starr and Fridolfsson, 2010). Plant LINC complexes contain conserved SUN proteins but have a divergent complement of KASH proteins (Graumann et al., 2010; Zhou et al., 2012). The WIP and WIT protein families form a LINC complex with Arabidopsis SUN1 and SUN2. This complex binds the Myosin XI-i motor protein and is involved in moving nuclei in root and leaf cells (Tamura et al., 2013). In pollen, the WIT-WIP-SUN complex is involved in male fertility. Loss of either the WIP or WIT protein family, as well as loss of both, results in a 50% reduction in seed set and a reversed order within the MGU upon emergence from the pollen grain (Zhou and Meier, 2014). While pollen tube growth is not significantly altered, the MGU distance from the growing pollen tube tip is increased (Zhou and Meier, 2014). Frequently, the VN is not observed at the pollen tube tip, and the corresponding pollen tubes either stall at the entrance to an ovule or continue to grow past the synergids but remain intact (Leydon et al., 2013; Zhou and Meier, 2014). How the requirement for the WIT-WIP-SUN complex, and, by extension, the tip-located VN, relates to the known steps of pollen tube reception and termination signaling is not known.

Here, we hypothesized that the WIT-WIP-SUN LINC complex is required for ROS-induced pollen tube rupture. We report a decrease in Ca^2+^-dependent ROS-induced pollen tube rupture of *wip* and *wit* mutants grown semi-*in vivo*. The hyposensitivity to ROS correlates with an increased distance of the VN from the pollen tube tip. In addition, we provide a first report of nuclear Ca^2+^ spiking in a gametophytic nucleus and show the patterns during growth and ROS-mediated pollen tube burst are altered in the mutants.

## Results and Discussion

This study uses two previously described Arabidopsis mutant lines “*wit12”* (T-DNA insertions in *WIT1* and *WIT2*) and “*wifi”* (T-DNA insertions in *WIP1*, *WIP2*, *WIP3*, *WIT1*, and *WIT2*) that have defects in MGU trafficking and pollen tube rupture (Zhou and Meier, 2014). Pollen tubes were tested for their sensitivity to external application of hydrogen peroxide (H_2_O_2_) by performing *semi-in vivo* pollen tube rupture assays at 5 hours post pollination (hpp) and at 7 hpp, as described previously (Duan et al., 2014). Pollen tube rupture events were defined by the appearance of cytoplasmic material at the end of the pollen tube (Fig. 1A, B), as described before (Duan et al., 2014). At 5 hpp, approximately 45% of Columbia-0 wild type (WT) pollen tubes treated with H_2_O_2_ burst, corroborating prior studies (Duan et al., 2014). In contrast, a reduced number of rupture events was observed in *wit12* and *wifi* pollen, with a burst frequency of 37% and 34% respectively (Fig. 1C).

**Figure 1.**
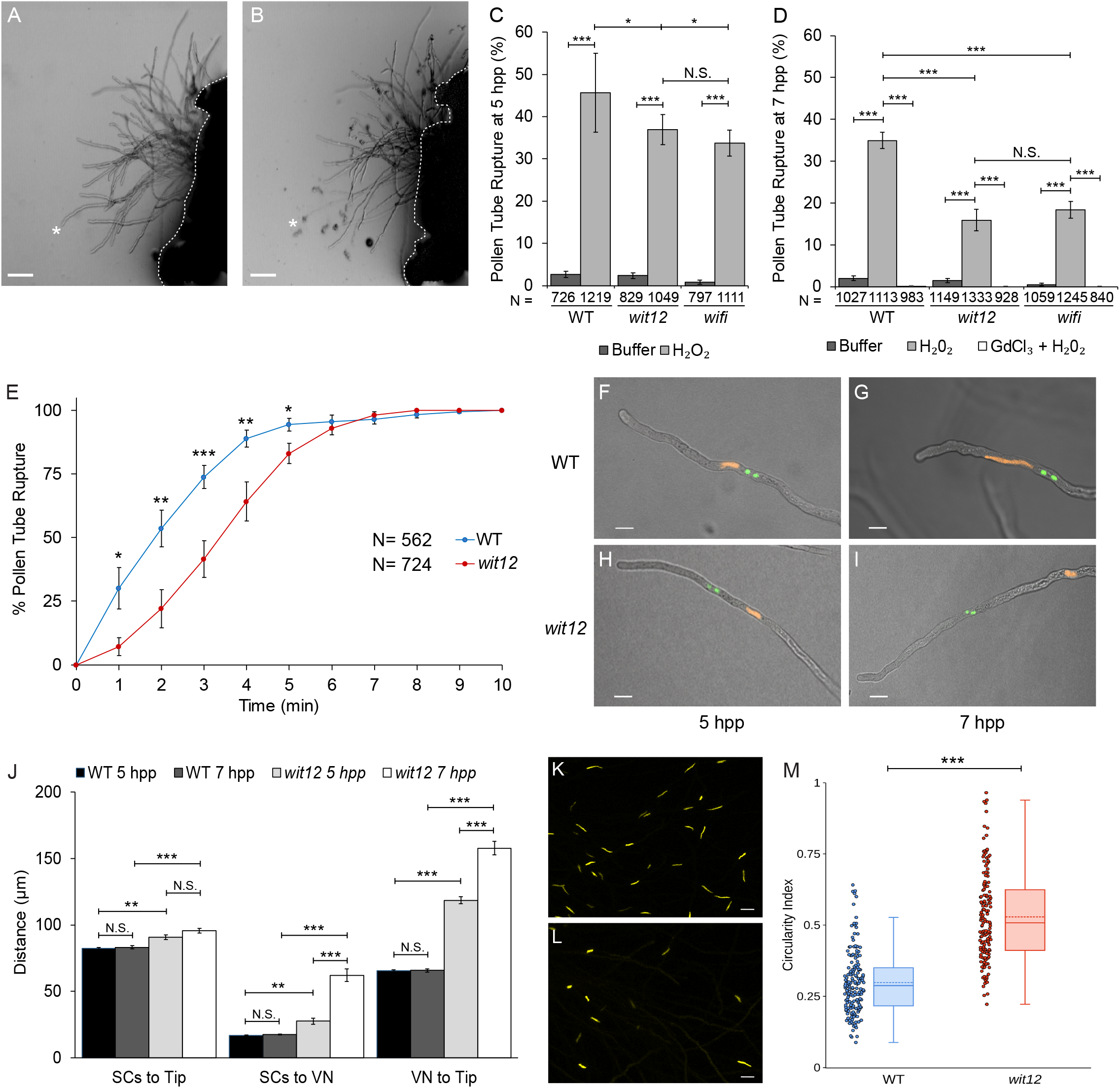
Ca^2+^-dependent ROS-induced pollen tube burst correlates with nuclear position. **A-B**, Semi*-in vivo* germinated WT pollen expressing *Lat52*_*pro*_*::R-GECO1* Ca^2+^ sensor. Size bars = 50µm. **A**, Pollen tubes prior to incubation with the ROS generating compound H_2_O_2_. **B**, Pollen tubes following incubation with H_2_O_2_. Several tubes have ruptured as indicated by cytoplasm outside the pollen tube. The asterisks in A and B mark the same pollen tube before and after rupture. The white dotted line marks the base of the stigma. **C-D**, ROS induced pollen tube rupture in semi-*in vivo* germinated pollen at 5 hours (**C**) and 7 hours (**D**) post pollination (hpp). Values represent the average number of ruptured pollen tubes observed when treated with buffer, H_2_O_2_, or H_2_O_2_ and a Ca^2+^ channel inhibitor, gadolinium(III) chloride (GdCl_3_ + H_2_O_2_). Bars are standard error. N equals total number of pollen tubes. * P≤0.05; **P≤0.01; *** P≤0.001 for Student’s t-test; N.S. indicates no significance by Student’s t-test. **E**, The time to ROS induced pollen tube rupture in semi-*in vivo* germinated WT and *wit12* pollen at 7 hpp. Bars are standard error. N equals total number of pollen tubes. * P≤0.05; **P≤0.01; *** P≤0.001 for Student’s t-test. **F-I**, The localization of the VN (orange) and SCs (green) was examined in elongating pollen tubes at 5 and 7 hpp. The VN and SCs were visualized using *Ub10*_*pro*_::*NLS-mCherry* and *MGH3*_*pro*_::*GFP* respectively. The position of the VN tip in WT remains close to the pollen tube tip at 5 (**F**) and 7 hpp (**G**). The VN in *wit12* pollen increases in distance from the pollen tube tip over time. The nucleus is closer at 5 hpp (**H**) than at 7 hpp (**I**). Scale bars = 10µm. **J**, The position of the SCs and VN relative to the tip and distance between the SCs and VN were quantified for WT and *wit12* pollen tubes at both time points. N = 150 pollen tubes. Bars are standard error. **P≤0.01; *** P≤0.001 for Student’s t-test. **K-L**, Representative images of a WT VN (**K**) and a *wit12* VN (**L**) at 7 hpp, using the YFP signal of NLS-YC3.6 as a nuclear marker. **M**, Circularity index of vegetative nuclei from WT and *wit12* pollen tubes. (Left) Scatter plot showing each nucleus as a single point. (Right) Box plot. (Top line) Maximum. (Box) Quartiles. (Solid middle line) Median. (Dotted middle line) Mean. (Bottom line) Bottom fence. ***P<0.001 for Student’s t-test.

When the experiment was extended by another 2 hours (7 hpp), efficiency of burst dropped in WT pollen tubes to 35% but dropped substantially more in *wit12* (16%) and *wifi* (18%). At both 5 hpp and 7 hpp, the two mutants behaved similarly (Fig. 1C, D). This suggests that *semi-in vivo* pollen tube burst in *wit12* and *wifi* is hyposensitive to external ROS, and that this effect increases over time. When pollen tubes were grown on low calcium (Ca^2+^) media and treated with the Ca^2+^ channel inhibitor gadolinium(III) chloride prior to the application of H_2_O_2_, all three lines behaved similarly with few to no rupture events observed (Figure 1D). In addition, we analyzed time to response of pollen tubes rupture within 10 minutes after addition of H_2_O_2_ (Figure 1E). The data show that there was an overall delay in response in the *wit12* pollen tube population compared to WT.

Together, these data show that *wit12* and *wifi* are similarly hyposensitive to ROS-induced pollen tube rupture, that this hyposensitivity increases over time post pollination, and that while pollen tubes burst less frequently in *wit12* and *wifi*, the burst still depends on Ca^2+^.

Because mutations in *WIP* and *WIT* affect the mobility of the VN during pollen tube growth, we tested whether the nuclear position plays a role in ROS response. Because *wit12* and *wifi* pollen tubes showed a comparable response to ROS, as well as comparable male fertility defects (Zhou and Meier, 2014), we proceeded with the *wit12* mutant only. WT and *wit12* pollen tubes concurrently expressing the VN (*Ub10pro::NLS-mCherry*) and SC (*MGH3pro::GFP*) fluorescent markers were germinated using the semi-*in vivo* method and the nuclear position was examined. In WT, the VN remained at a constant distance of approximately 65 µm from the pollen tube tip, regardless of when the pollen tubes were examined (Figure 1F, 1G). By contrast, at 5 hpp the VN of *wit12* pollen tubes was positioned, on average, 118 µm from the pollen tube tip (Figure 1H, 1J). This distance was further increased to an average of 157 µm at 7 hpp (Figure 1I, 1J). The separation between VN and SCs in *wit12* also increased between 5 hpp and 7 hpp (Figure 1J).

In addition, we noted that *wit12* nuclei appeared less elongated than WT nuclei, which were thin and highly stretched (Figure 1K, 1M). Circularity index was calculated for WT and *wit12* nuclei, to determine if a quantifiable difference in circularity exists. When compared to WT, *wit12* nuclei are shorter and more circular (Figure 1L, 1M). Taken together, these data indicate a correlation between VN-tip distance and responsiveness to exogenous ROS in the pollen tube rupture mechanism.

A relationship between position of the nucleus and the strength of a signaling pathway has been shown in animals, for example in the notch signaling pathway (Del Bene et al., 2008). In this case, a diffusible compound is involved and the signal reaching the nucleus is dampened by increased distance. Reception of the signal requires a LINC complex for nuclear movement, a shared similarity with the *wit12* and *wifi* mutants described here (Del Bene et al., 2008; Zhou and Meier, 2014). In Arabidopsis pollen, mutants in a family of transcription factors (MYB97/101/120) show similar pollen tube reception defects as *wit12* and *wifi* mutants (Leydon et al., 2013; Liang et al., 2013). A diffusible signal might thus be upstream of the VN gene expression profile required for late stages of pollen tube growth.

As an obvious candidate for such a signal, we tested whether the cytoplasmic Ca^2+^ signatures of growing pollen tube tips were impaired. Transgenic lines expressing the cytoplasmic Ca^2+^sensor R-GECO1 under the pollen-specific promoter Late Anther Tomato 52 (LAT52) (Eyal et al., 1995) were used in time-lapse imaging of WT and *wit12* pollen tubes at 7 hpp (Figure 2A, 2B). Transgenic pollen was germinated using the semi-*in vivo* method. The fluorescence signatures were measured at the tips of growing WT and *wit12* pollen tubes using a region of interest (ROI) based analysis method in the NIS Elements analysis software (see Materials and Methods). Qualitatively, both WT and *wit12* pollen tubes exhibited a dynamic fluorescence pattern (Figure 2C, 2D) consistent with what has been previously reported for growing Arabidopsis pollen tubes (Iwano et al., 2009). Variability in observed fluorescence intensities between WT and *wit12* may be due to different insertional sites between transgenic lines. To determine differences in Ca^2+^ fluctuations between WT and *wit12*, the traces were quantified in two ways, by counting the number of peaks after setting a threshold value (Figure 2E), and by calculating the average standard deviation of all traces (Figure 2F) (see Materials and Methods). WT and *wit12* exhibited similar peak numbers, with an average of 7 peaks within 10 minutes of imaging (Figure 2E). The standard deviation between WT and *wit12* was also similar (Figure 2F). These data suggest that the cytoplasmic Ca^2+^ signatures at the pollen tube tip of *wit12* are not significantly altered during elongation.

**Figure 2.**
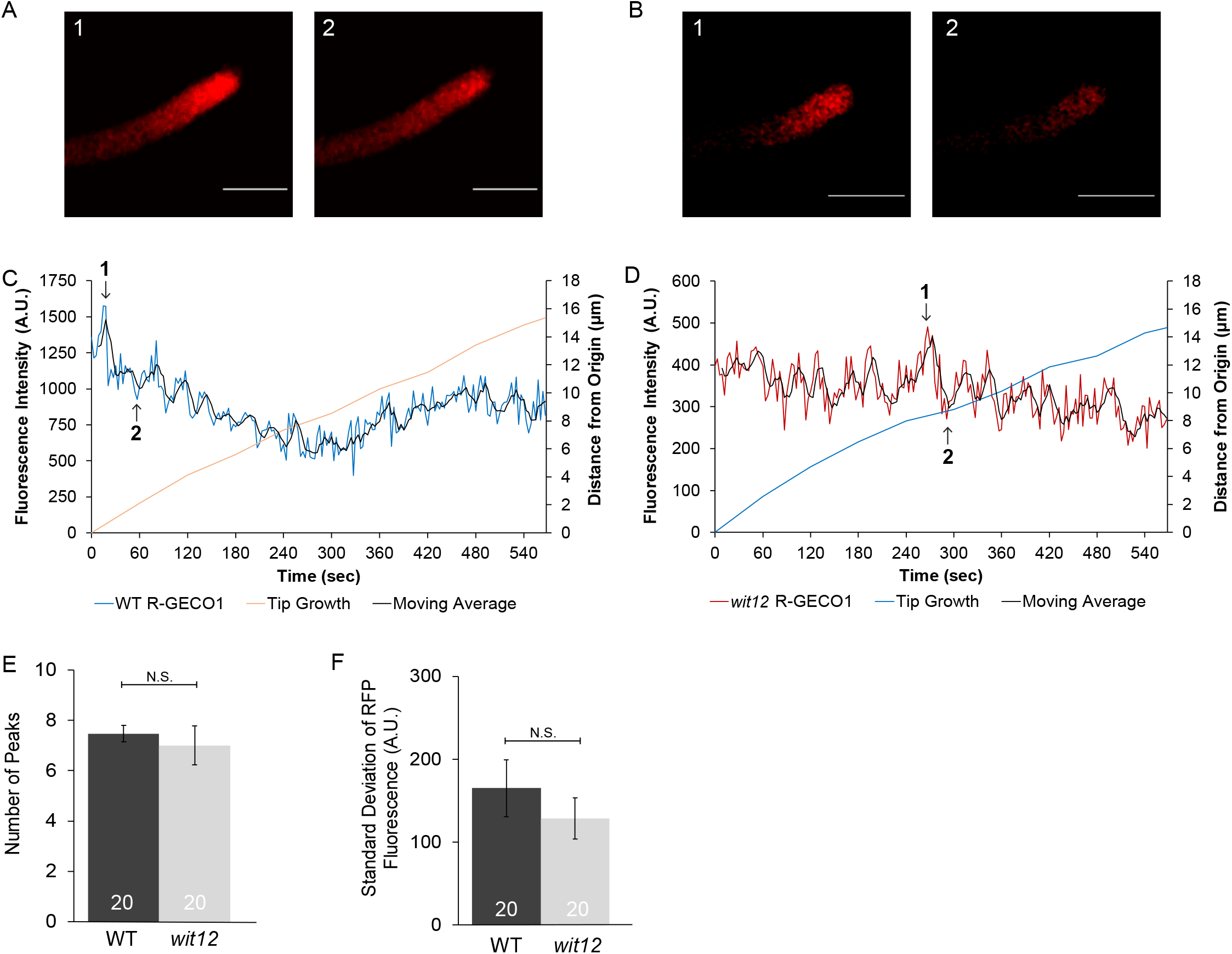
Cytoplasmic Ca^2+^ fluctuations are not disrupted in *wit12* pollen tubes. **A-F**, The cytoplasmic Ca^2+^ sensor R-GECO1 (*Lat52*_*pro*_::*R-GECO1*) was used to measure Ca^2+^ signatures in elongating pollen tubes. **A-B**, Images of changes in cytoplasmic Ca^2+^ fluctuations during growth for a WT pollen tube (**A**) and a *wit12* pollen tube (**B**). Images in **A** and **B** correspond to time points 1 and 2, shown in **C** and **D**. Scale bar = 10µm. **C**, Cytoplasmic Ca^2+^ fluctuations measured at the pollen tube tip of WT pollen are represented as fluorescence intensity in the blue line. The red line marks the pollen tube growth from the origin at the start of imaging. The black line represents a 4-point rolling average of the data used to highlight the overall trend. Numbers in the graph correspond to the image numbers in **A**. **D**, Cytoplasmic Ca^2+^ fluctuations at the tip of elongating *wit12* pollen. Fluorescence intensity is depicted by the red line. The blue line marks the pollen tube growth from the origin at the start of imaging. The black line represents a 4-point rolling average of the data used to highlight the overall trend. Numbers in the graph correspond to the image numbers in **B**. **E-F**, Cytoplasmic Ca^2+^ signatures were quantified based on the number of peaks (**E**) and the standard deviation of the fluorescence intensity for the entire time-lapse movie (**F**). 20 pollen tubes were analyzed in each case. Bars are standard error. N.S. indicates no significant difference by Student’s t-test.

A change in the Ca^2+^-dependent capacity for ROS-induced rupture that correlates with nuclear position might suggest a nuclear Ca^2+^ signaling step. Several examples for nuclear Ca^2+^ signaling have been reported in plants, ranging from plant-microbe interactions to development (Capoen et al., 2011; Charpentier et al., 2016; Leitão et al., 2019). To observe nuclear Ca^2+^, we generated transgenic WT and *wit12* plants containing the VN-localized Ca^2+^ sensor NLS-YC3.6, driven by the LAT52 promoter (Figure 3A-3C). WT nuclei exhibited clear Ca^2+^ fluctuations at 7 hpp, with an average of 3.5 peaks within 10 minutes of imaging (Figure 3D, 3F). In contrast, the Ca^2+^ signature observed in elongating *wit12* pollen tubes exhibited one of two distinct patterns. In the first case, the number of Ca^2+^ fluctuations were reminiscent of those observed in WT (Figure 3E). In the second case, the number of Ca^2+^ fluctuations were reduced compared to WT or no fluctuations were observed (Figure 3E).

**Figure 3.**
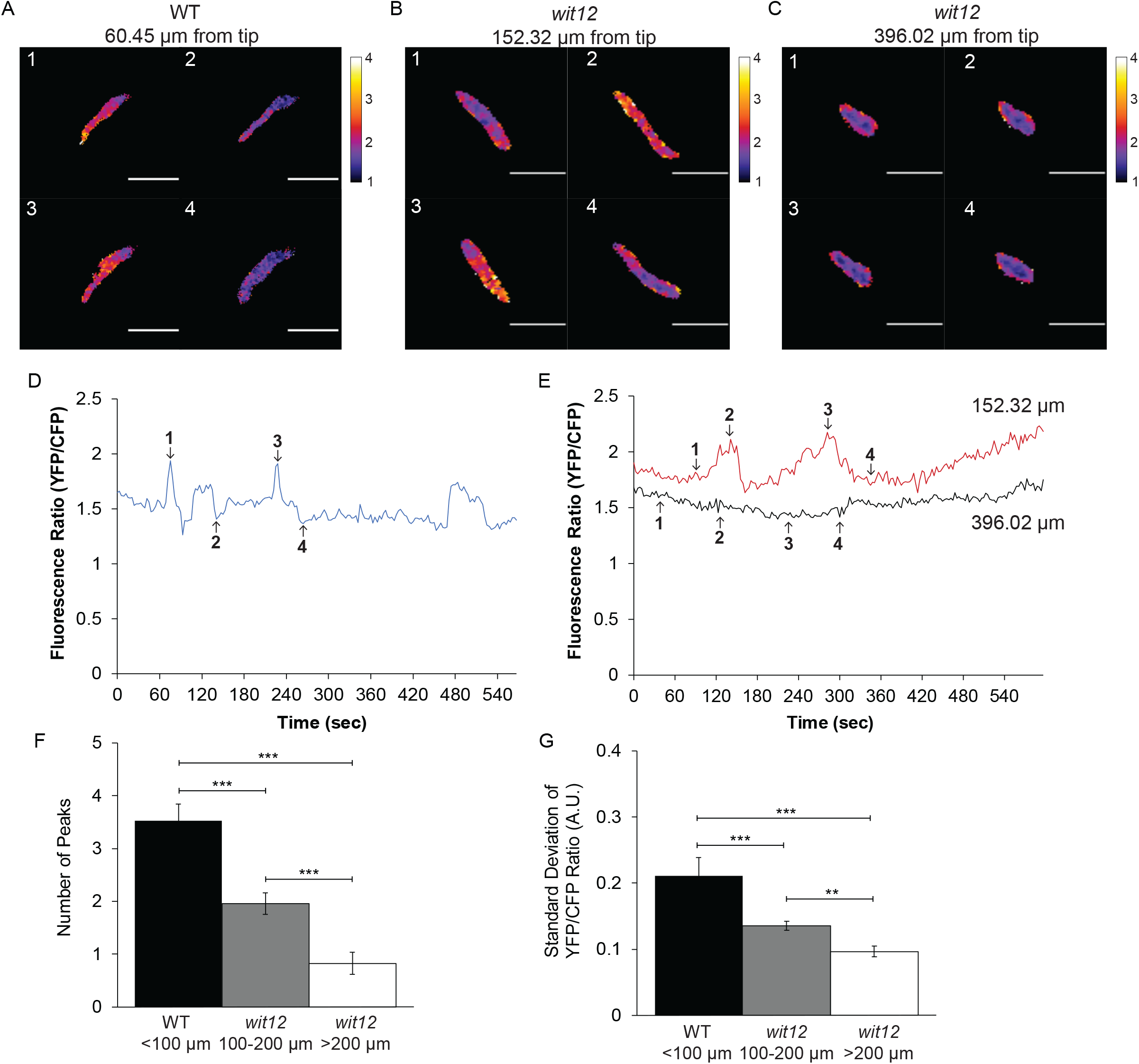
Pollen nuclear Ca^2+^ signatures correlate with the position of the nucleus. **A-G**, The nucleus-localized Ca^2+^ sensor (*Lat52*_*pro*_::NLS-YC3.6) was used to measure Ca^2+^ signatures in the VNs of elongating pollen tubes **A-C**, Ratio images of changes in nuclear Ca^2+^ fluctuations during growth for a WT pollen tube with the VN positioned 60 µm from the tip (**A**), a *wit12* pollen tube with the VN positioned 152 µm from the tip (**B**), a *wit12* pollen tube with the VN positioned 396 µm from the tip (**C**). Scale bar = 10µm. **D**, A representative nuclear Ca^2+^ signature for a WT nucleus 60 µm from the pollen tube tip, presented as FRET ratio of YFP to CFP fluorescence intensity, is shown as a blue line. Numbers in the graph correspond to the image numbers in **A**. **E**, A representative nuclear Ca^2+^ signature for a *wit12* nucleus 152 µm from the pollen tube tip, presented as a FRET ratio of YFP to CFP fluorescence intensity, is shown as a red line. A representative nuclear Ca^2+^ signature for a *wit12* nucleus 396 µm from then pollen tube tip, presented as a FRET ratio of YFP to CFP fluorescence intensity, is shown as a black line. Numbers in the graph correspond to the image numbers in **B** and **C**, respectively. **F-G**, Nuclear Ca^2+^ signatures were quantified based on the number of peaks (**F**) and the standard deviation of the YFP/CFP ratio for the entire time-lapse movie (**G**). At least 20 pollen tubes were analyzed per background. Columns were split based on the distance of the VN from the pollen tube tip. Bars are standard error. * P<0.05; **P<0.01; *** P<0.001.

We next tested if the reduction in nuclear Ca^2+^ fluctuations observed in *wit12* was dependent on the distance of the VN to the pollen tube tip. For each 10-minute movie generated, the distance between the VN and tip was also recorded. The WT nuclei were positioned between 50 and 100 µm from the tip (Figure 3F, 3G), consistent with data presented in Figure 1J. VNs in *wit12* were distributed over a range of tip distances, but never below 100 µm. *wit12* VNs located between 100 µm and 200 µm from the pollen tube tip exhibited a more WT-like Ca^2+^ signature, although the average frequency of peaks per 10 minutes was reduced to 2 peaks (Figure 3F, 3G). *wit12* VNs located further than 200 µm from the pollen tube tip exhibited a more disrupted Ca^2+^ signature, on average showing only 1 peak (Figure 3F, 3G). These data show that there are nuclear Ca^2+^ fluctuations during pollen tube growth and that the frequency of fluctuations is correlated with the proximity of the VN to the pollen tube tip.

Nuclear Ca^2+^ fluctuations have been described as compounds of signal transduction pathways in a variety of biological systems, notably during root symbioses (Capoen et al., 2011; Liang et al., 2014; Charpentier et al., 2016). After nodulation (Nod) or mycorrhizal (Myc) factor perception, pronounced Ca^2+^ oscillations in and around the nucleus are observed, which are required for the transcriptional response of the host plant (Charpentier, 2018). Examples of nuclear Ca^2+^ spiking were also found in relation to biotic and abiotic stress as well as root development; however, in these cases single or few spikes were observed instead of pronounced oscillations (Charpentier, 2018; Leitão et al., 2019).

The Ca^2+^ signal signatures observed here are similar to those observed after biotic or abiotic stress (Lecourieux et al., 2005; Kelner et al., 2018). This response is dampened in the *wit12* mutant. Furthermore, *wit12* mutant pollen tubes showing the greatest reduction in Ca^2+^ spiking have the greatest distance between VN and pollen tube tip. It has been shown that plant nuclei can generate their own Ca^2+^ signals, independent of changes in cytoplasmic Ca^2+^, and that a simple increase of the extranuclear Ca^2+^ concentration is not sufficient to trigger Ca^2+^ increases in isolated nuclei (Pauly et al., 2000; Mazars et al., 2009). It is therefore unlikely that the nuclear Ca^2+^ spikes in the pollen tube simply reflect cytoplasmic Ca^2+^ fluctuations by passive Ca^2+^ influx. Instead, a cytoplasmic Ca^2+^ signal could be required to directly or indirectly activate a nuclear Ca^2+^ signal (Charpentier, 2018), and distance from the tip could therefore dampen this response.

Finally, we examined how the cytoplasmic and nuclear Ca^2+^ signatures changed in ^response to H_2_O_2_^. WT pollen tubes that ruptured exhibited first an increase in the cytoplasmic calcium signal at the apex of the pollen tube (Figure 4A, top row). The increase in signal then propagated down the shank while the signal at the apex remained elevated (Figure 4A). These observations are in agreement with what was described previously (Duan et al., 2014). In *wit12* pollen tubes, a similar sequence of events was observed prior to burst, suggesting that there is no significant difference in the H_2_O_2_-induced tip Ca^2+^ response (Figure 4A, bottom row).

**Figure 4.**
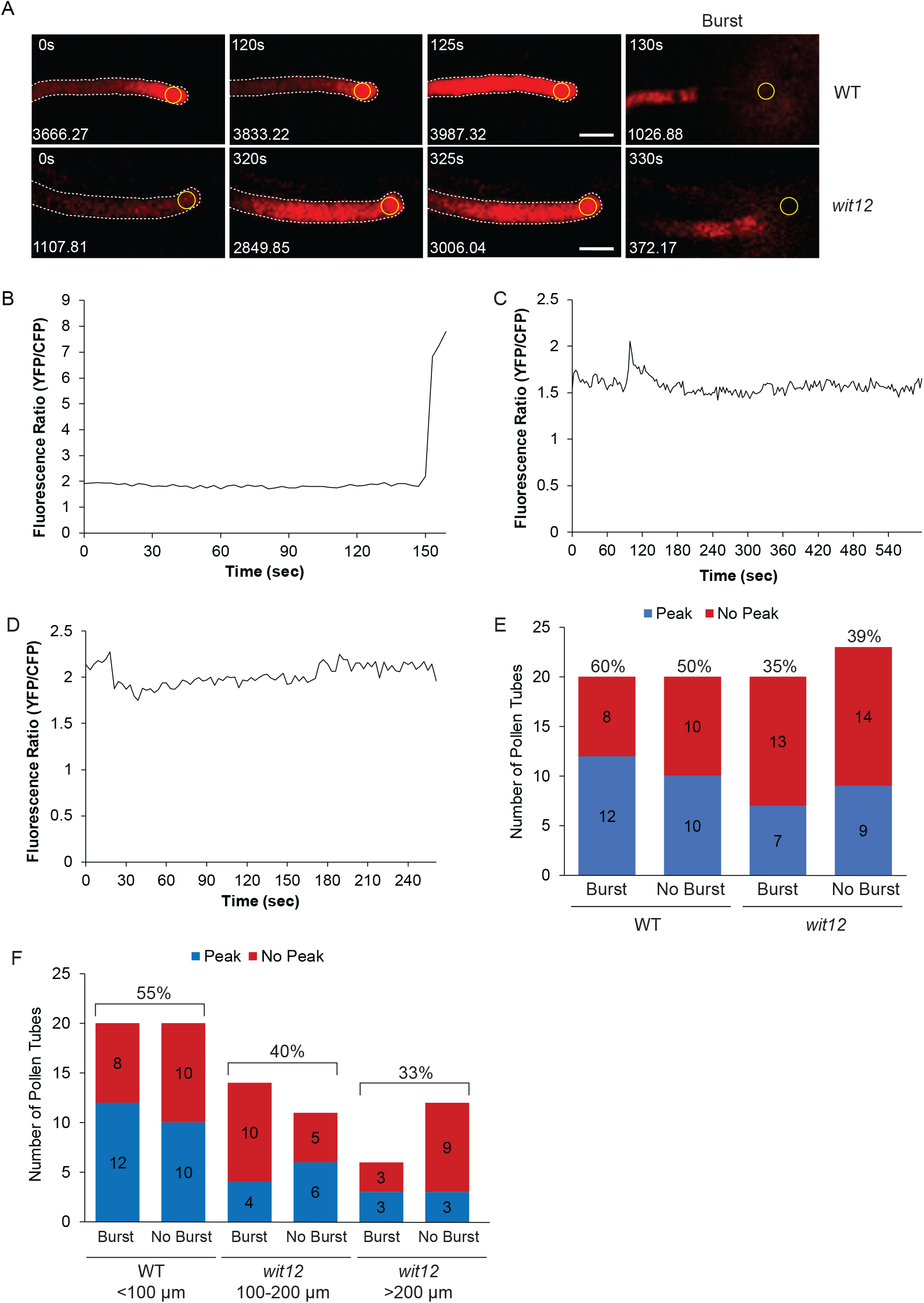
Post-ROS Ca^2+^ fluctuations. **A**, The Ca^2+^ sensor R-GECO1 (*Lat52*_*pro*_::*R-GECO1*) was used to measure cytoplasmic Ca^2+^ signatures over time in pollen tubes after addition of H_2_O_2_. Representative images of changes in cytoplasmic Ca^2+^ fluctuations for one WT pollen tube (top panel) and one *wit12* pollen tube (bottom panel). The white dotted line outlines the pollen tube wall using the corresponding DIC image. Numbers in the bottom left corner are the RFP fluorescence of the corresponding ROIs (yellow circles), located close to the pollen tube tip. Time 0 indicates the start of imaging after H_2_O_2_ addition. Imaging sequences shown are representative of several videos for each genotype. Scale bar = 10 µm. **B-F**, The Ca^2+^ sensor *Lat52*_*pro*_::NLS-YC3.6 was used to measure nuclear Ca^2+^ signatures over time in the VNs after addition of H_2_O_2_. **B-D**, Examples of the different types of nuclear Ca^2+^ signatures observed in WT after addition of H_2_O_2_. A range of nuclear Ca^2+^ spikes are shown, from the highest peak height of 7 A.U. (**B**) to the lowest peak height of 0.5 A.U. (**C**). **D**, A nuclear Ca^2+^ signature without a spike. **E-F**, Quantification of presence or absence of peaks for pollen tubes that burst and pollen tubes that fail to burst for both WT and *wit12*. The numbers of pollen tubes with or without a peak for each condition is shown in the bar graph. **E**, WT is compared to the entire *wit12* population. The percent of pollen tubes that presented a nuclear Ca^2+^ peak is shown above each bar. **F**, *wit12* columns were split based on the distance of the VN from the pollen tube tip. The percent of pollen tubes that presented a nuclear Ca^2+^ peak, with burst and no burst combined for each group, is shown above each bar.

Next, we described the Ca^2+^ signature in the VN of WT and *wit12* pollen in response to H_2_O_2_. Pollen tubes, expressing the NLS-YC3.6 sensor, were treated with H_2_O_2_ at 7 hpp. A Ca^2+^ peak was defined as a signal that both increased and decreased at least 0.5 A.U., or that increased at least 0.5 A.U. immediately prior to rupture. A range of nuclear Ca^2+^ peaks were recorded, from a peak height of 7 A.U. to the lowest peak counted of 0.5 A.U. (Figures 4B, 4C). Within both populations, we observed pollen tubes that showed a peak either immediately prior to rupture (Figure 4B), a peak followed by a signal decline (Figure 4C), or no Ca^2+^ peak as defined here (Figure 4D). 60% of WT pollen tubes that burst showed a peak, compared to 50% that did not burst (Figure 4E). 35% of *wit12* pollen tubes that burst and 39% of *wit12* pollen tubes that did not burst showed a peak (Figure 4E). Thus, overall fewer *wit12* pollen tubes showed a post-H_2_O_2_ nuclear Ca^2+^ spike compared to WT, but this reduction was not correlated with whether the pollen tubes burst or did not burst.

Regardless of whether the pollen tube ruptured, 55% of WT pollen tubes had a Ca^2+^ spike (Figure 4F). When the *wit12* population was split up based on the distance of the VN from the pollen tube tip, 40% of VNs positioned between 100 µm and 200 µm from the tip showed a Ca^2+^ spike, compared to only 33% VNs located over 200 µm from the tip (Figure 4F). In addition, of the 20 pollen tubes randomly chosen that did burst, 14 had the VN between 100 µm and 200 µm and only 6 over 200 µm from the tip. Among the 23 non-bursting *wit12* pollen tubes, the VN was over 200 µm from the tip in 12 cases, again suggesting that the capacity to undergo H_2_O_2_-induced burst correlates with the position of the nucleus.

Together, these data suggest that there is a correlation between nuclear Ca^2+^ spiking and VN position, as well as between H_2_O_2_-induced burst and VN position, but there is no correlation between nuclear Ca^2+^ spiking in individual pollen tubes and the capacity of the specific pollen tube to rupture.

The relationship at the single-cell level between the VN Ca^2+^ spiking and ROS-induced pollen tube burst remains unclear. Nevertheless, this is the first evidence for nuclear Ca^2+^ spikes in a higher plant gametophyte, and in light of the known relevance of nuclear Ca^2+^ signaling in the sporophyte, likely functionally relevant. It is possible that the VN senses cytoplasmic Ca^2+^ signatures, including those elevated by ROS, but that the nuclear Ca^2+^ spikes trigger downstream signals unrelated to pollen tube burst.

Newly developed dual Ca^2+^ sensors (Kelner et al., 2018) will be better equipped to resolve the timing relationship between cytoplasmic and nuclear Ca^2+^ signatures and the relationship between the spread of cytoplasmic Ca^2+^ in the shank and the position of the nucleus. Discovering and investigating male gametophyte-expressed nuclear envelope-associated Ca^2+^ channels can now address which processes are affected by disrupting VN Ca^2+^ signatures.

## Supplemental Data

The following materials are available in the online version of this article.

**Supplemental Table S1: Primers used for cloning.** CACC sites for directional TOPO cloning are indicated in bold. Specific recognition sites for *SacI* and *SpeI* are underlined.

**Supplemental Movie S1**

An example of cytoplasmic Ca^2+^ signatures in a WT pollen tube after addition of H_2_O_2_. The Ca^2+^ sensor R-GECO1 (*Lat52*_*pro*_*::R-GECO1)* was used to measure changes in Ca^2+^ over time. The pollen tube tip was imaged every 5 seconds until rupture. Movie playback speed is 60x.

**Supplemental Movie S2**

An example of cytoplasmic Ca^2+^ signatures in a *wit12* pollen tube after addition of H_2_O_2_. The Ca^2+^ sensor R-GECO1 (*Lat52*_*pro*_*::R-GECO1)* was used to measure changes in Ca^2+^ over time. The pollen tube tip was imaged every 5 seconds until rupture. Movie playback speed is 60x.

## Acknowledgements

This work was funded by a National Science Foundation grant to I.M. (NSF-1613501). We would like to thank all members of the Meier lab for many fruitful discussions throughout this work. We thank Dr. Keith Slotkin (Donald Danforth Plant Science Center) for gifting the *LAT52*_*pro*_::*GFP* construct, Dr. Melanie Krebs (Universität Heidelberg) for gifting *UBQ10*_*pro*_::*NLS-YC3.6*, Dr. Simon Gilroy (University of Wisconsin-Madison) for gifting *35S*_*pro*_::*YC3.6* and Dr. Anna Dobritsa (The Ohio State University) for critically reading this manuscript.

## Materials and Methods

### Plant materials

*Arabidopsis thaliana* (Columbia-0 ecotype) was germinated on Murashige and Skoog medium plates (Caisson Laboratories) containing 1% sucrose under constant light. Plants at the two-leaf stage were transplanted to soil and grown at an average temperature of 22-23°C under a 16-hour light/8-hour dark regime. The *wit1-1* (GABI-Kat 470E06) *wit2-1* (SALK_127765) (*wit12*) double null mutant was reported by (Zhao et al., 2008) and the quintuple null mutant *wip1-1* (SAIL_390_A08) *wip2-1* (SALK_052226) *wip3-1* (GABI-Kat 459H07) *wit1-1 wit2-1* (“*wifi”*) mutant was reported by (Zhou et al., 2012). Heterozygous *male sterility-1* (*ms-1*) plants were obtained from the Arabidopsis Biological Resource Center (https://abrc.osu.edu/).

### Cloning

All primers used in cloning and construct generation are outlined in Supplemental Table 1. The pollen specific promoter LAT52 was cloned from the *LAT52*_*pro*_::*GFP* construct in the binary plasmid pMDC107 as previously reported (Zhou and Meier, 2014). Restriction sites for enzymes *Sac*I and *Spe*I were added to the 5’ and 3’ ends respectively. The amplified fragment was digested with the appropriate restriction enzymes. The LAT52 promoter fragment was isolated, purified with the QIAquik PCR Purification kit (Qiagen) and subsequently ligated into a pH2GW7 vector (Takagi et al., 2011).

The Yellow Cameleon 3.6 (YC3.6) calcium (Ca^2+^) sensor N-terminally tagged with a nuclear localization signal (NLS) was cloned from a *UBQ10*_*pro*_::*NLS-YC3.6* (Krebs et al., 2011). The R-GECO1 Ca^2+^ sensor construct was amplified from *CMV*_*pro*_*::R-GECO1* (Zhao et al., 2011). NLS-YC3.6 and R-GECO1 were cloned into pENTR/D-TOPO vectors (Life Technologies) and then moved to *LAT52*_*pro*_::*pH2GW7* via the Gateway® LR reaction (Life Technologies).

### Generation of transgenic plants

Binary vectors containing Ca^2+^ sensors were transformed into either *Agrobacterium tumefaciens* strains ABI or GV3101 by triparental mating (Wise et al., 2006). Arabidopsis plants were transformed using the *Agrobacterium*-mediated floral dip method (Clough and Bent, 1998). The Col-0 ecotype (WT), *wit12* and *wifi* backgrounds were used for the transformation. Transgenic plants were isolated on MS plates supplemented with using 30 µg/mL hygromycin, and positive transformants were confirmed by confocal microscopy. Homozygous T2 plants were used for all assays described here.

### Semi *in-vivo* pollen germination and ROS-induced pollen tube rupture

Pollen germination media (PGM) containing 5mM KCl, 5mM CaCl_2_, 1 mM Ca(NO_3_)_2_, 1mM MgSO_4_, 10% sucrose, 0.01% boric acid, pH 7.5 (Palanivelu and Preuss, 2006) was heated with 0.4% (w/v) agarose, pipetted onto a glass slide, and allowed to solidify before use. Anthers at time of anthesis were used to pollinate stigmas of *male sterility 1* (*ms-1*) flowers at developmental stage 14 (Zhou and Meier, 2014). Two hours after pollination, stigmas were excised and placed horizontally onto the solid PGM agar pad. The stigmas were incubated for an additional 3 or 5 hours in a humidity chamber (Palanivelu and Preuss, 2006). Elongating pollen tubes were imaged using a 4X objective with 5 times digital zoom. At time of imaging, pollen tubes were treated with 10µL of liquid PGM (control) or with 10µL of liquid PGM containing 6mM H_2_O_2_.

Immediately following treatment, single plane images of pollen tubes were acquired every 10 seconds for 10 minutes with confocal microscopy (Eclipse C90i; Nikon). In addition, after this 10-minute time course, images were acquired at different focal planes to ensure that all pollen tubes could be identified and counted. To quantify the percentage of rupture events, the total number of intact and germinated pollen tubes were tabulated prior to and 10 minutes after treatment. The number of pollen tube discharge events were then counted. For the Ca^2+^ depletion experiments, pollen was germinated on low-Ca^2+^ PGM medium (5mM KCl, 100µM CaCl_2_, 1mM MgSO_4_, 10% sucrose, 0.01% boric acid, pH 7.5). Pollen tubes were then treated with 10µL of low-Ca^2+^ PGM containing 100 µM GdCl_3_ and after a 10-minute incubation 10µL of low-Ca^2+^ PGM with 12mM H_2_O_2_ was added.

### Time to rupture quantification

The time to rupture was quantified using data collected in Figure 1D. For each experiment, the 10-minute videos were split into 1-minute intervals and the number of pollen tubes that burst during each interval was recorded. That number was then divided by the total number of pollen tubes that ruptured over the course of the 10-minute video and represented as percent. The average percent rupture for each time point was plotted.

### Measurement of positions of male germ unit components within the pollen tube

WT and *wit12* lines expressing both the VN marker *Ub10*_*pro*_::*NLS-mCherry* and SC marker *MGH3*_*pro*_::*GFP* were germinated using the semi-*in vivo* method on PGM agar pads, as described above. Elongated pollen tubes were imaged using a 40X water immersion objective. Images of pollen at 5 and 7 hpp were acquired at several different focal planes to ensure the MGU and pollen tube tip were in focus. Distances from the relative center of the sperm cells (SCs) and vegetative nucleus (VN) to the tip of the pollen tube were measured using the NIS Elements analysis software (Nikon).

### Ca^2+^ sensor imaging

Pollen expressing either the R-GECO1 Ca^2+^ sensor or the NLS-YC3.6 nuclear Ca^2+^ sensor was germinated as previously described (Boavida and McCormick, 2007). A square was drawn onto a glass slide with a grease pencil (Staples). An isolated, pollinated stigma was placed in the center of the square. The square was filled with liquid PGM until the media began to mound. The slide was placed in a high humidity chamber for 5 hours to allow pollen tube elongation (Boavida and McCormick, 2007; Johnson and Kost, 2010). Individual elongating pollen tubes were imaged under a coverslip with a 40X water immersion objective with 2 x digital zoom. NLS-YC3.6 sensor was excited with 457 nm, and fluorescence emission was detected between 465 and 505 nm (CFP) and between 530 nm and 570 (cpVenus). R-GECO1 was excited with 561 nm, and its emission was detected between 620 and 650 nm. Time lapse movies of the pollen tube tip and VN were generated by acquiring an image every 3 seconds for a total of 10 minutes. Because expression levels were highly variable between transgenic lines and pollen tubes, laser and gain settings were adjusted individually to obtain comparable baseline intensity values for each experiment. To quantify the Ca^2+^ signature, a region of interest (ROI) was defined using the binary editor function of the NIS Elements analysis software. For cytoplasmic Ca^2+^, an ROI was drawn proximal to the pollen tube tip. For nuclear Ca^2+^, an ROI was drawn around the VN. The mean fluorescence intensity was obtained for the defined ROI for every image captured in the time-lapse dataset and graphed as a function of time (“Calcium Signature”). For NLS-YC3.6, the YFP/CFP ratio for each time point was calculated by dividing the mean fluorescence intensity of YFP by the mean fluorescence intensity of CFP.

### Peak number and standard deviation measurement

Ca^2+^ signatures during pollen tube growth were quantified based on the number of peaks and the standard deviation of the RFP signal for R-GECO1 and the YFP/CFP ratio for NLS-YC3.6. Cytoplasmic peaks were determined based on an RFP fluorescence intensity increase of 150 A.U. or higher from the baseline, while nuclear Ca^2+^ peaks were determined based on a YFP/CFP ratio increase of 0.25 A.U. or higher. The average fluorescence intensity or YFP/CFP ratio and the standard deviation for each 200-frame time-lapse movie was calculated. All 20 standard deviations were then averaged to determine the mean standard deviation for R-GECO1 in WT, R-GECO1 in *wit12*, NLS-YC3.6 in WT, and NLS-YC3.6 in *wit12*. Nuclear Ca^2+^ signatures after addition of H_2_O_2_ were analyzed using a similar peak determination as described above. However, a nuclear Ca^2+^ peak was determined based on a YFP/CFP ratio increase of 0.50 A.U. or higher.

### Circularity index measurement

WT and *wit12* pollen tubes expressing *Lat52*_*pro*_::NLS-YC3.6 were grown using the semi-*in vivo* method and imaged at 7 hpp with a confocal microscope (Eclipse C90i, Nikon). Images were taken using a 40X water immersion objective and acquired using NIS-Elements AR version 3.2. At least 150 pollen vegetative nuclei were visualized for each line by creating z-stack sections (3 μm) to capture the entire nucleus. A maximum intensity projection of each nucleus was generated, and ImageJ was used to calculate the circularity index of each nucleus.

### Accession Numbers

Sequence data from this article can be found in the GenBank/EMBL data libraries under accession numbers NM_001160797 (WIP1), NM_125004 (WIP2), NM_112181 (WIP3), NM_121177 (WIT1), and NM_105565 (WIT2).

## References

Boavida LC, McCormick S (2007) Temperature as a determinant factor for increased and reproducible in vitro pollen germination in Arabidopsis thaliana. Plant J 52: 570–582

Boisson-Dernier A, Roy S, Kritsas K, Grobei MA, Jaciubek M, Schroeder JI, Grossniklaus U (2009) Disruption of the pollen-expressed FERONIA homologs ANXUR1 and ANXUR2 triggers pollen tube discharge. Development 136: 3279–3288

Borg M, Brownfield L, Twell D (2009) Male gametophyte development: a molecular perspective. J Exp Bot 60: 1465–1478

Capoen W, Sun J, Wysham D, Otegui MS, Venkateshwaran M, Hirsch S, Miwa H, Downie JA, Morris RJ, Ané J-M, Oldroyd GED (2011) Nuclear membranes control symbiotic calcium signaling of legumes. Proc Natl Acad Sci USA 108: 14348

Charpentier M (2018) Calcium signals in the plant nucleus: origin and function. J Exp Bot 69: 4165–4173

Charpentier M, Sun J, Martins TV, Radhakrishnan GV, Findlay K, Soumpourou E, Thouin J, Véry A-A, Sanders D, Morris RJ, Oldroyd GED (2016) Nuclear-localized cyclic nucleotide–gated channels mediate symbiotic calcium oscillations. Science 352: 1102

Clough SJ, Bent AF (1998) Floral dip: a simplified method for Agrobacterium-mediated transformation of Arabidopsis thaliana. Plant J 16: 735–743

Del Bene F, Wehman AM, Link BA, Baier H (2008) Regulation of neurogenesis by interkinetic nuclear migration through an apical-basal notch gradient. Cell 134: 1055–1065

Dresselhaus T, Franklin-Tong N (2013) Male-female crosstalk during pollen germination, tube growth and guidance, and double fertilization. Mol Plant 6: 1018–1036

Duan Q, Kita D, Johnson EA, Aggarwal M, Gates L, Wu H-M, Cheung AY (2014) Reactive oxygen species mediate pollen tube rupture to release sperm for fertilization in Arabidopsis. Nat Commun 5: 3129

Dumas C, Knox R, Gaude T (1985) The spatial association of the sperm cells and vegetative nucleus in the pollen grain of *Brassica*. Protoplasma 124: 168–174

Escobar-Restrepo J-M, Huck N, Kessler S, Gagliardini V, Gheyselinck J, Yang W-C, Grossniklaus U (2007) The FERONIA Receptor-like Kinase Mediates Male-Female Interactions During Pollen Tube Reception. Science 317: 656

Eyal Y, Curie C, McCormick S (1995) Pollen specificity elements reside in 30 bp of the proximal promoters of two pollen-expressed genes. Plant Cell 7: 373–384

Ge Z, Bergonci T, Zhao Y, Zou Y, Du S, Liu MC, Luo X, Ruan H, García-Valencia LE, Zhong S, Hou S, Huang Q, Lai L, Moura DS, Gu H, Dong J, Wu HM, Dresselhaus T, Xiao J, Cheung AY, Qu LJ (2017) pollen tube integrity and sperm release are regulated by RALF-mediated signaling. Science 358: 1596–1600

Graumann K, Runions J, Evans DE (2010) Characterization of SUN-domain proteins at the higher plant nuclear envelope. Plant J 61: 134–144

Higashiyama T, Yang W-c (2017) Gametophytic Pollen Tube Guidance: Attractant Peptides, Gametic Controls, and Receptors. Plant Physiol 173: 112

Iwano M, Entani T, Shiba H, Kakita M, Nagai T, Mizuno H, Miyawaki A, Shoji T, Kubo K, Isogai A, Takayama S (2009) Fine-Tuning of the Cytoplasmic Ca(2+) Concentration Is Essential for Pollen Tube Growth. Plant Physiol 150: 1322–1334

Iwano M, Ngo QA, Entani T, Shiba H, Nagai T, Miyawaki A, Isogai A, Grossniklaus U, Takayama S (2012) Cytoplasmic Ca2+ changes dynamically during the interaction of the pollen tube with synergid cells. Development 139: 4202–4209

Johnson MA, Harper JF, Palanivelu R (2019) A Fruitful Journey: Pollen Tube Navigation from Germination to Fertilization. Annu Rev Plant Biol 70: 809–837

Johnson MA, Kost B (2010) Pollen Tube Development. Methods Mol Biol 655: 155–176

Kelner A, Leitão N, Chabaud M, Charpentier M, de Carvalho-Niebel F (2018) Dual Color Sensors for Simultaneous Analysis of Calcium Signal Dynamics in the Nuclear and Cytoplasmic Compartments of Plant Cells. Front Plant Sci 9: 245

Kessler SA, Grossniklaus U (2011) She’s the boss: signaling in pollen tube reception. Curr Opin Plant Biol 14: 622–627

Krebs M, Held K, Binder A, Hashimoto K, Den Herder G, Parniske M, Kudla J, Schumacher K (2011) FRET-based genetically encoded sensors allow high-resolution live cell imaging of Ca2+ dynamics. Plant J 69: 181–192

Lecourieux D, Lamotte O, Bourque S, Wendehenne D, Mazars C, Ranjeva R, Pugin A (2005) Proteinaceous and oligosaccharidic elicitors induce different calcium signatures in the nucleus of tobacco cells. Cell Calcium 38: 527–538

Leitão N, Dangeville P, Carter R, Charpentier M (2019) Nuclear calcium signatures are associated with root development. Nat Commun 10: 4865

Leydon AR, Beale KM, Woroniecka K, Castner E, Chen J, Horgan C, Palanivelu R, Johnson MA (2013) Three MYB transcription factors control pollen tube differentiation required for sperm release. Curr Biol 23: 1209–1214

Li H-J, Meng J-G, Yang W-C (2018) Multilayered signaling pathways for pollen tube growth and guidance. Plant Reprod 31: 31–41

Liang Y, Tan ZM, Zhu L, Niu QK, Zhou JJ, Li M, Chen LQ, Zhang XQ, Ye D (2013) MYB97, MYB101 and MYB120 function as male factors that control pollen tube-synergid interaction in Arabidopsis thaliana fertilization. PLoS Genet 9: e1003933

Liang Y, Tóth K, Cao Y, Tanaka K, Espinoza C, Stacey G (2014) Lipochitooligosaccharide recognition: an ancient story. New Phytol 204: 289–296

Mazars C, Bourque S, Mithöfer A, Pugin A, Ranjeva R (2009) Calcium homeostasis in plant cell nuclei. New Phytol 181: 261–274

McCue AD, Cresti M, Feijó JA, Slotkin RK (2011) Cytoplasmic connection of sperm cells to the pollen vegetative cell nucleus: potential roles of the male germ unit revisited. J Exp Bot 62: 1621–1631

Ngo QA, Vogler H, Lituiev DS, Nestorova A, Grossniklaus U (2014) A calcium dialog mediated by the FERONIA signal transduction pathway controls plant sperm delivery. Dev Cell 29: 491–500

Palanivelu R, Preuss D (2006) Distinct short-range ovule signals attract or repel Arabidopsis thaliana pollen tubes in vitro. BMC Plant Biol 6: 7

Pauly N, Knight MR, Thuleau P, van der Luit AH, Moreau M, Trewavas AJ, Ranjeva R, Mazars C (2000) Control of free calcium in plant cell nuclei. Nature 405: 754–755

Russell S, Cass D (1981) Ultrastructure of the sperms of *Plumbago zeylanica* 1. Cytology and association with the vegetative nucleus. Protoplasma 107: 85–107

Starr DA, Fridolfsson HN (2010) Interactions between nuclei and the cytoskeleton are mediated by SUN-KASH nuclear-envelope bridges. Annu Rev Cell Dev Biol 26: 421–444

Takagi S, Islam MS, Iwabuchi K (2011) Dynamic Behavior of Double-Membrane-Bounded Organelles in Plant Cells. Int Rev Cell Mol Biol 286: 181–222

Tamura K, Iwabuchi K, Fukao Y, Kondo M, Okamoto K, Ueda H, Nishimura M, Hara-Nishimura I (2013) Myosin XI-i Links the Nuclear Membrane to the Cytoskeleton to Control Nuclear Movement and Shape in Arabidopsis. Curr Biol 23: 1776–1781

Wise AA, Liu Z, Binns AN (2006) Three Methods for the Introduction of Foreign DNA into Agrobacterium. Methods Mol Biol 343: 43–54

Zhang J, Huang Q, Zhong S, Bleckmann A, Huang J, Guo X, Lin Q, Gu H, Dong J, Dresselhaus T, Qu LJ (2017) Sperm cells are passive cargo of the pollen tube in plant fertilization. Nat Plants 3: 17079

Zhao Q, Brkljacic J, Meier I (2008) Two distinct interacting classes of nuclear envelope-associated coiled-coil proteins are required for the tissue-specific nuclear envelope targeting of Arabidopsis RanGAP. Plant Cell 20: 1639–1651

Zhao Y, Araki S, Wu J, Teramoto T, Chang YF, Nakano M, Abdelfattah AS, Fujiwara M, Ishihara T, Nagai T, Campbell RE (2011) An expanded palette of genetically encoded Ca^2+^ indicators. Science 333: 1888–1891

Zhou X, Graumann K, Evans DE, Meier I (2012) Novel plant SUN-KASH bridges are involved in RanGAP anchoring and nuclear shape determination. J Cell Biol 196: 203–211

Zhou X, Meier I (2014) Efficient plant male fertility depends on vegetative nuclear movement mediated by two families of plant outer nuclear membrane proteins. Proc Natl Acad Sci USA 111: 11900–11905

